# The impact of chemical fixation on the microanatomy of mouse brain tissue

**DOI:** 10.1101/2023.02.21.528828

**Authors:** Agata Idziak, V.V.G. Krishna Inavalli, Stephane Bancelin, Misa Arizono, U. Valentin Nägerl

## Abstract

Chemical fixation using paraformaldehyde (PFA) is a standard step for preserving cells and tissues for subsequent microscopic analyses such as immunofluorescence or electron microscopy. However, chemical fixation may introduce physical alterations in the spatial arrangement of cellular proteins, organelles and membranes. With the increasing use of super-resolution microscopy to visualize cellular structures with nanometric precision, assessing potential artifacts - and knowing how to avoid them - takes on special urgency.

We addressed this issue by taking advantage of live-cell super-resolution microscopy that makes it possible to directly observe the acute effects of PFA on organotypic brain slices, allowing us to compare tissue integrity in a ‘before-and-after’ experiment. We applied super-resolution shadow imaging to assess the structure of the extracellular space (ECS) and regular super-resolution microscopy of fluorescently labeled neurons and astrocytes to quantify key neuroanatomical parameters.

While the ECS volume fraction and micro-anatomical organization of astrocytes remained largely unaffected by the PFA treatment, we detected subtle changes in dendritic spine morphology and observed substantial damage to cell membranes. Our experiments show that PFA application via immersion does not cause a noticeable shrinkage of the ECS in brain slices, unlike the situation in transcardially perfused animals where the ECS typically becomes nearly depleted.

In addition to the super-resolved characterization of fixation artefacts in identified cellular and tissue compartments, our study outlines an experimental strategy to evaluate the quality and pitfalls of various fixation protocols for the molecular and morphological preservation of cells and tissues.

## Introduction

Chemical fixation is a commonly used preservation step for electron microscopy (EM) and super-resolution microscopy techniques, such as Stimulated Emission Depletion microscopy (STED), single-molecule localization microscopy and expansion microscopy. These techniques permit structural and molecular analyses of cells and tissues at a sub-microscopic level. Chemical fixatives like paraformaldehyde (PFA) covalently cross-link proteins, which has the effect of physically hardening the cellular and molecular structure of the sample. This procedure is to protect the sample from decay and damage during subsequent processing steps, such as tissue slicing, dehydration or embedding in resin.

However, it is known that even the most carefully executed fixation protocol may introduce structural artifacts that compromise data quality and interpretation (Ebersold et al., 1981; Maugel and Hayat, 1977; Ryter, 1988; Schnell et al., 2012). While these problems may not be noticeable at a macroscopic level, they can appear at the microscopic subcellular scale. Indeed, organelles such as endosomes and lysosomes become deformed by chemical fixation (Murk et al., 2003), while cellular proteins can still move substantially and reposition after chemical fixation, potentially casting doubts over conclusions based on this approach (Tanaka et al., 2010).

As the spatial resolution of microscopy techniques keeps improving, allowing researchers to make ever more detailed and discriminating observations, concerns about fixation artifacts become more relevant. Recent super-resolution techniques can now reach into the low nanometer range, where fixation artifacts may abound. In turn, these gains in spatial resolution necessitate the development of more stringent ways to assess the quality of fixation protocols and how well they can preserve cellular elements at this finer spatial scale.

The question of how much the micro-architecture and ultrastructure of brain tissue is affected by chemical fixation was addressed in two EM studies that compared the effects of chemical and cryogenic fixation protocols on tissue fine structure. Cryogenic fixation is based on rapid high-pressure freezing of the sample, which produces amorphous ice instead of ice crystals, that otherwise would destroy the ultrastructure. These studies clearly showed that chemical fixation via transcardial perfusion leads to a strong depletion of the extracellular space (ECS) as well as changes in astrocytic (Korogod et al., 2015) and dendritic spine morphology (Tamada et al., 2020), raising serious concerns about the use of chemical fixation protocols in high-resolution anatomical studies of brain tissue. However, due to differences in sample preparation required for either fixation method and the inability to compare the EM samples with their live originals, the reason of the observed differences remains elusive.

To directly compare nanoscale neuroanatomical structures before and after chemical fixation, we took advantage of the super-resolution shadow imaging (SUSHI) technique, which combines 3D-STED microscopy and fluorescence labeling of the interstitial fluid (Tønnesen et al., 2018). SUSHI allows for visualization of tissue anatomy, including the ECS, projecting all cellular structures as sharply contoured ‘shadows’, providing a comprehensive and non-biased view of the tissue. Using this technique, we imaged organotypic brain slices and analyzed the impact of PFA on the ECS. In addition, we imaged fluorescently labeled astrocytes and neurons, and analyzed the effect of PFA on their nanoscale morphology in a before-and-after manner.

We observed that PFA does not induce major changes in the shape and size of the ECS and astrocytes. However, we detected subtle changes in dendritic spine morphology as well as a widespread disruption of cellular membranes and cellular blebbing.

The study gives a ‘real time’ and nanoscale view of the effects of PFA on brain tissue micro-architecture, revealing the extent and type of fixation artifacts, which had remained inconclusive. The super-resolution approach based on positive and inverse labeling provides an accurate and comprehensive readout of the impact of chemical fixation on brain tissue, facilitating the optimization of fixation protocols to preserve the native structure of the tissue as well as possible.

## Results

### 30 minutes of PFA fixation has no noticeable effects on ECS volume fraction

To directly assess whether chemical fixation using PFA has an effect on hippocampal ECS structure, we established an experimental workflow that allowed us to compare the same sample before and after PFA fixation in a paired manner (**Fig. 1A**).

**Figure 1.**
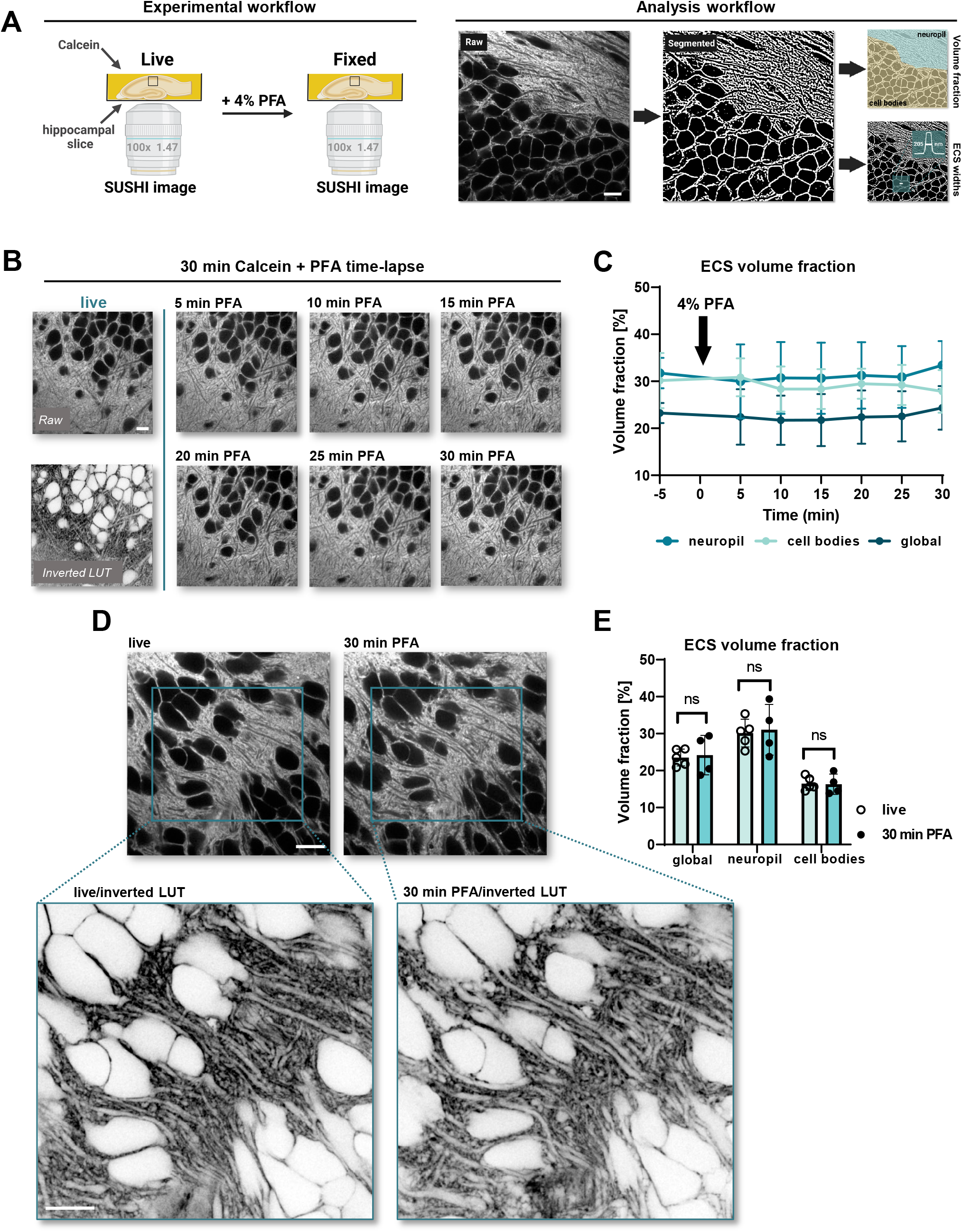
Brief PFA application does not affect ECS volume fraction. **(A)** Graphical overview of the workflow of experiments and analysis. **(B)** Time-lapse shadow imaging of ECS in living and PFA-fixed conditions. The live condition is represented both with a raw and inverted LUT. **(C)** ECS volume fraction changes over 30 minutes of PFA fixation. The images were analyzed either as a whole (‘global’) or divided into ‘neuropil’ or ‘cell bodies’ areas (n_**cntrl**_ = 5; n_**PFA**_ = 6; ns: not significant, *P < 0.05; in paired student t-test). **(D)** Representative images of ECS live and 30 minutes PFA-fixed. Blue squares indicate the magnified area, that is showed below in an inverted LUT. **(E)** Paired analysis of ECS volume fraction live and after 30 minutes of PFA fixation (n=5; ns: not significant, *P < 0.05; in Wilcoxon matched-pairs test). Scale bars: 10 µm.

We performed time-lapse confocal shadow imaging at 5-minute intervals before and during PFA application using Calcein to label the artificial cerebrospinal fluid (ACSF) that the slices were maintained in (**Fig. 1B**). The images were binarized using SpineJ software (Levet et al., 2020) based on wavelet filtering to calculate the ECS volume fraction (VF), which in our case was the ratio of the ECS area over the total area in a region of interest. We found that 30 min of PFA incubation did not cause any significant changes in ECS volume fraction (**Fig. 1C**; n_cntrl_ = 5, n_PFA_ = 6; p > 0.05, paired student t-test).

This result was confirmed by SUSHI (**Fig. 1D & E**; n = 6, p > 0.05; Wilcoxon matched-pairs test), indicating that 30 minutes of PFA incubation has little impact on the VF of the ECS in organotypic hippocampal slices.

### Prolonged PFA incubations introduce pronounced artifacts

As brain slices are often maintained in fixative for more than one hour or even overnight, we investigated the effects of longer incubation times on ECS VF (**Fig. 2A**). We imaged for 90 min under PFA conditions as well in regular ACSF, PFA-free conditions for control. While 90 minutes PFA fixation neither affected the ECS VF (**Fig. 2B left**; n = 6; p > 0.05, Wilcoxon matched-pairs test) nor ECS widths measured in line profiles of segmented images using SpineJ (**Fig. 2C, D, E;** n_cntrl_ = 12 lines, n_PFA_ = 16 lines; p_cntrol_ = 0.1281, P_pfa_ = 0.7249, paired t-test), we observed dye-free, cellular blebs in the immediate vicinity of cell bodies (**Fig 2A white arrow; Fig 2B right**).

**Figure 2.**
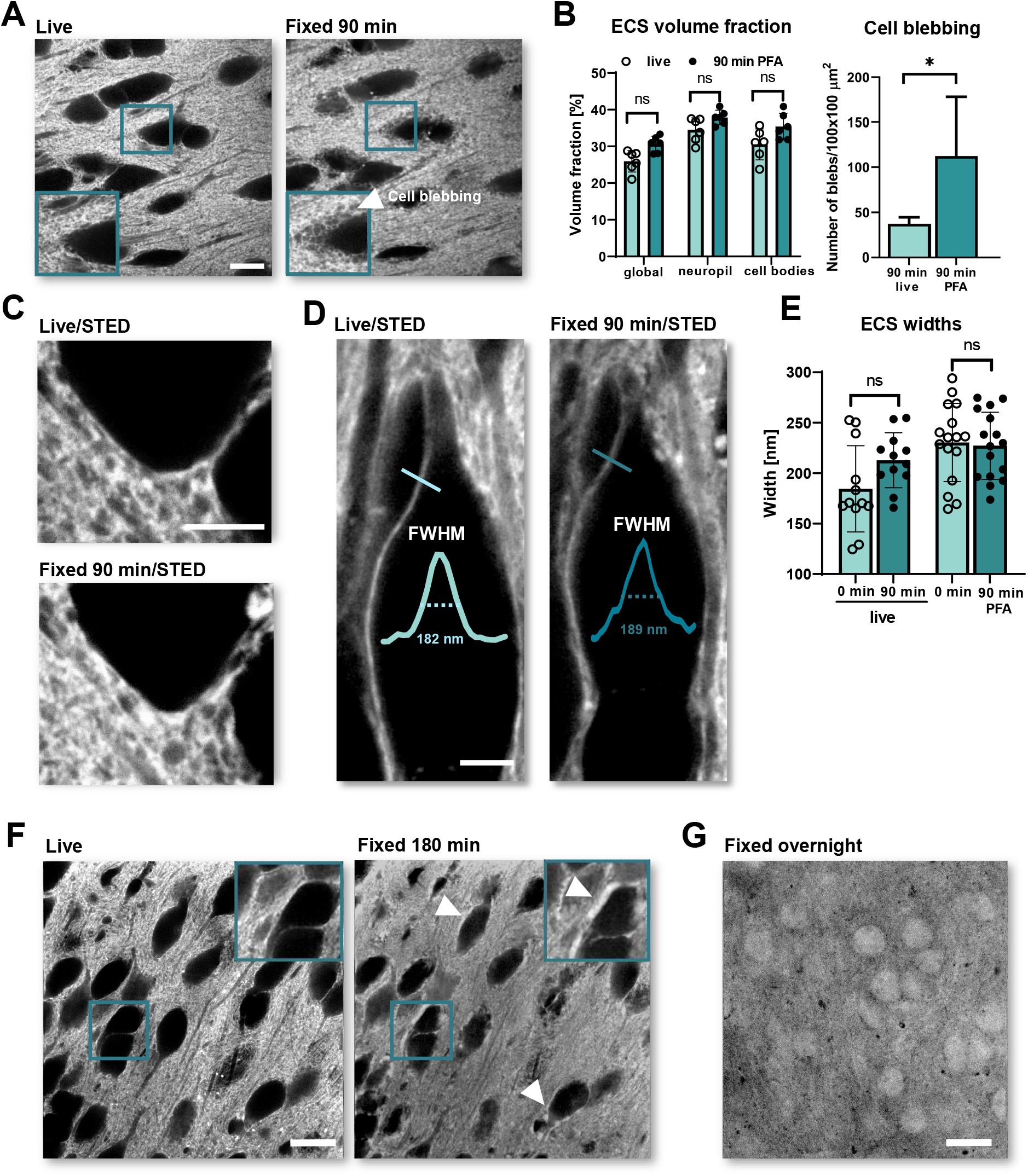
Prolonged PFA application introduces ECS artifacts. **(A)** Representative images of ECS live and 90 minutes after fixation. The inset and white arrow indicate ‘cell blebbing’ artefact. **(B) Left:** Paired analysis of ECS volume fraction between live and 90 minutes after PFA fixation. The images were analyzed either as a whole (‘global’) or divided into ‘neuropil’ or ‘cell bodies’ areas (n_**cntrl**_ = 6; n_**PFA**_ = 6; ns: not significant, *P < 0.05; in Wilcoxon matched-pairs test). **Right:** Comparison of the ‘cell blebbing’ between live and 90 minutes PFA-fixed conditions (n=6; ns: not significant, *P < 0.05; in a paired student t-test). **(C)** Representative STED images of ECS live and 90 minutes after PFA fixation. **(D)** SUSHI-based ECS width. The blue lines indicate an example of the analyzed width. The line profiles are shown together with a measured FWHMs. **(E)** Paired analysis of ECS widths live and 90 minutes after PFA fixation (n_**cntrl**_=12; n_**PFA**_=16; ns: not significant, *P < 0.05; in a paired t-test). **(F)** Representative images of ECS live and 180 minutes after PFA fixation. Inset and white arrows indicate examples of dye accumulation around cell bodies. **(G)** A representative image of shadow imaging after overnight fixation with PFA. Scale bars (A, F, G): 10 µm; (C, D): 5 µm.

180 minutes of PFA incubation caused even more prominent changes, such as dye accumulation around cell bodies and dye permeation into the cells (**Fig. 2F**), indicating that PFA incubation by itself permeabilized cell membranes, even in the absence of detergents, like Triton, that are typically used in immunofluorescence protocols to get antibodies to reach intracellular epitopes. Indeed, after overnight PFA incubation the extracellular dye had strongly penetrated into the cells (**Fig. 2G**), indicating disruption of cellular membranes. This made assessing the impact of PFA on ECS impossible, because of the loss of inside-outside contrast required for the shadow imaging approach. Thus, more than 90 minutes of PFA fixation appears to seriously damage the integrity of cellular membranes.

### 90 minutes of PFA fixation does not affect the morphology of astrocytes

Beside the effect on ECS volume and widths, we set out to determine the impact of PFA fixation on different cell types. We first focused on astrocytes, whose morphology is known to be very sensitive to environmental changes, such as osmotic challenges (Arizono et al., 2021), or transcardial perfusion (Korogod et al., 2015). In order to label astrocytes, we micro-injected AAV-GFAP-Clover viral particles into organotypic hippocampal slices. Confocal microscopy revealed no significant changes in the size of the major branches and cell bodies of astrocytes after 90 minutes of PFA fixation (**Fig. 3A, B**; n_branches_= 12; n_bodies_= 11; Wilcoxon matched-pairs test). Similarly, STED microscopy revealed no significant changes in the widths of fine astrocytic processes (**Fig. 3C, D**; n_cntrl_ = 26; n_PFA_= 28; Wilcoxon matched-pair test). These results suggest PFA incubation by itself has surprisingly little impact on astrocytic morphology.

**Figure 3.**
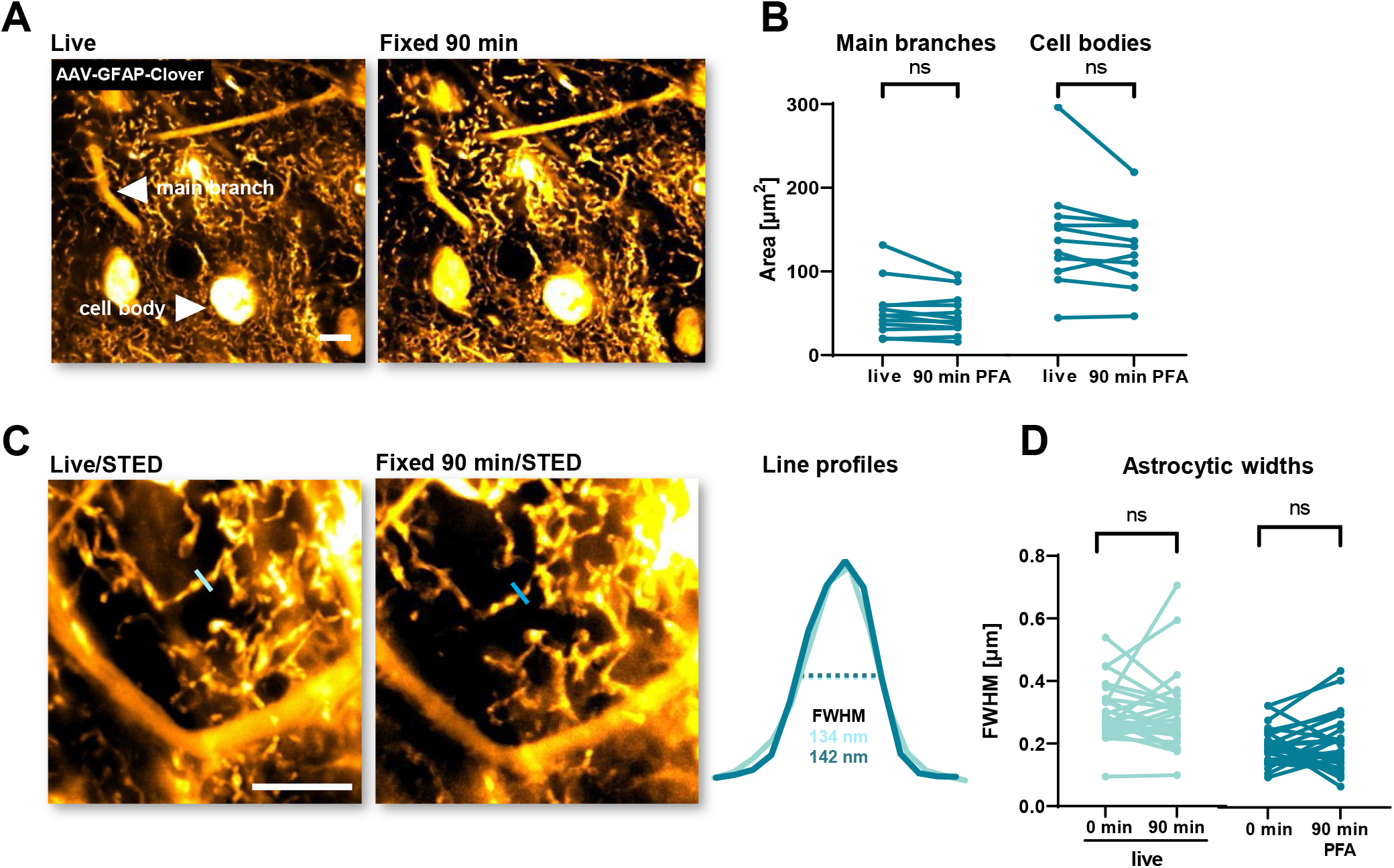
PFA fixation has no visible effects on nanoscale astrocytic morphology. **(A)** Representative confocal images of a brain slice expressing GFAP-Clover in astrocytes, live and 90 min after PFA fixation. White arrows indicate a representative astrocytic main branch and cell body analyzed. **(B)** Paired analysis of astrocytic areas of main branches and cell bodies live and 90 minutes after PFA fixation (n_**branches**_ = 12; n_**bodies**_ = 11; ns: not significant, *P < 0.05 in Wilcoxon matched-pairs test). **(C)** Representative STED images of astrocytic nanoscale spongiform structures expressing GFAP-Clover, live and 90 min after PFA fixation. The blue lines show a representative line across astrocytic structure for width analysis. Their profiles are shown on the right together with calculated FWHMs. **(D)** Paired analysis of astrocytic fine widths live and 90 minutes after PFA fixation (n_**cntrl**_ = 26; n_**PFA**_ = 28; ns: not significant, *P < 0.05; in Wilcoxon matched-pairs test). Scale bars: 10 µm.

### 90 minutes of PFA fixation leads to changes in dendritic spine morphology

Finally, we also performed similar experiments with neurons by virally labeling them with Citrine as fluorescent protein. Using STED microscopy, we imaged dendrites and dendritic spines before and after 90 minutes of PFA fixation (**Fig. 4A**). Unlike astrocytes, many dendrites formed ‘holes’ when exposed to 90 minutes of PFA, which could be genuine perforations in the dendritic membrane or pathological vacuoles free of the fluorescent label. Control experiments rule out the possibility that STED imaging was responsible for these artifacts (**Fig. 4B**; n_cntrl_ = 23, n_PFA_ = 31; p < 0.0001; one-sample Wilcoxon test).

**Figure 4.**
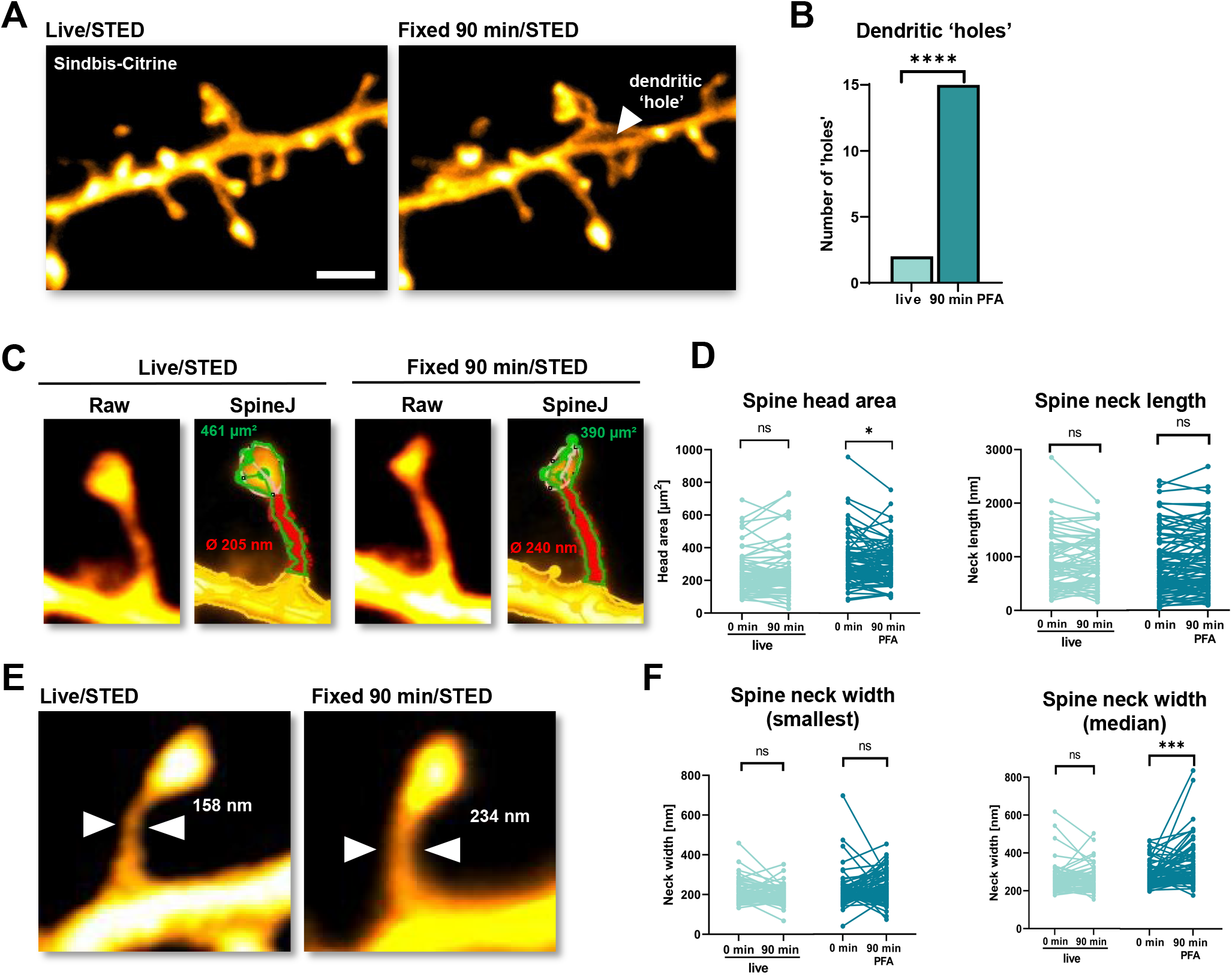
PFA fixation affects spine morphology. **(A)** Representative STED maximum intensity z-projections of a dendritic segment expressing cytosolic Citrine, live and 90 minutes after PFA fixation. A white arrow indicates a dendritic ‘hole’. Scale bar: 10 µm. **(B)** Bar graph showing the analysis of dendritic vacuoles appearing in live of 90 min of PFA fixation conditions (n_**cntrl**_ = 23; N_**pfa**_ = 31; one-sample Wilcoxon test). **(C)** Representative STED images of dendritic spines and an example of head and neck analysis using SpineJ. **(D)** Paired analysis of spine head area and neck length (n_**cntrl**_ = 79; n_**PFA**_ = 86; ns: not significant, *P < 0.05; in Wilcoxon matched-pairs test). **(E)** An example of a dendritic spine with a smaller neck width after 90 minutes of PFA fixation. **(F)** Paired analysis of spine neck width (smallest and median values) between live and 90 min PFA-fixed conditions (n_**cntrl**_ = 79; n_**PFA**_ = 86; ns: not significant, *P < 0.05; ***P < 0.001 in Wilcoxon matched-pairs test).

Detailed analysis of dendritic spine morphology revealed that there were no significant changes in spine neck lengths **(Fig. 4D)**; however, spine head area became significantly smaller **(Fig. 4C, D)** and the median spine neck diameter (measured along the length of the neck, see Methods for details) became wider after 90 minutes of PFA treatment (**Fig. 4E, F**; p_h.area_ = 0.0157; p_n.width_ < 0.0001; Wilcoxon matched-pair test). While this effect was highly statistically significant for the median neck diameter, it was not for the thinnest parts of the spine necks (**Fig. 4F**; p_n.width_ = 0.1565; Wilcoxon matched-pair test).

These results show that both dendrites and spines are affected by PFA fixation, suggesting that neurons are more sensitive than astrocytes to PFA treatment within the limits of the resolution of our nanoscale imaging approach.

## Discussion

In this study, we report the impact of PFA chemical fixation on brain tissue architecture, focusing on the ECS and cellular fine morphology. Our SUSHI approach revealed that simple immersion of organotypic brain slices in PFA has no appreciable effect on ECS volume fraction and widths. In the same vein, STED imaging showed no PFA-induced alterations in astrocytic morphology at the level of the soma, major branches and even their fine processes. At first sight, this is in contrast with a previous study that reported major ECS shrinkage or astrocytic swelling upon transcardial perfusion of PFA (Korogod et al., 2015). However, these pronounced changes may reflect an acute response *in vivo* to transcardial perfusion with PFA, such as anoxia (Tao-Cheng et al., 2007), and not a direct result of PFA by itself on the tissue. This is in line with the observation that acute slices fixed either chemically or cryogenically did not appear different in terms of tissue quality and ECS distribution (Korogod et al., 2015). Given the absence of major remodeling of the ECS under our conditions, it is not surprising perhaps that the astrocytes did not show any changes either, suggesting that their morphology and ECS topology are closely linked.

*In vivo* measurements have confirmed that conventional EM sample preparation drastically reduces ECS widths (Thorne and Nicholson, 2006). Due to slow speed of perfusion, transcardial fixation can also impact subcellular anatomy such as the spatial and molecular organization of synaptic vesicles (Maus et al., 2020). These structural changes in the synaptic environment (synapse, astrocytes and ECS) undoubtedly skew our understanding of synapse physiology, requiring improved protocols for *in vivo* fixation of brain tissue. In the same vein, a recent study showed that PFA can affect the behavior of proteins in liquid–liquid phase separation experiments, underscoring the importance of understanding better the artifacts induced by PFA fixation (Irgen-Gioro et al., 2022).

While having little impact on the morphology of astrocytes and ECS, PFA fixation over 90 minutes considerably disturbed membrane integrity as indicated by the penetration of the extracellular dye into the cells despite the arrestation of any active endocytic activity. This effect prevented us from performing SUSHI experiments because the contrast between the inside and outside of the cells disappeared. The membrane permeabilization was accompanied with cellular blebbing that has already been shown in cell cultures (Nanolive.ch, 2019; Zhao et al., 2014), suggesting that this is a common effect of PFA. Additional immunofluorescence experiments with and without membrane permeabilization confirm that PFA permeabilizes the membrane to an extent that a full-size antibody can pass into the cells (data not shown). Such loss of membrane integrity needs to be considered in experiments focused on membrane proteins, including ion channels or surface receptors (Ichikawa et al., 2022). In fact, many intracellular proteins are intimately linked to surface proteins via scaffold proteins forming large macro-molecular complexes, such as the postsynaptic density in dendritic spines (Chen et al., 2008). A loss of membrane integrity may distort our 3D spatial view of the synapse and the results of biophysical simulation studies based on protein localization obtained from fixed tissue.

While PFA fixation does not induce appreciable alterations in astrocytic structure, STED analysis of dendrites revealed that 90 minutes of PFA fixation results in dendritic ‘holes’. Their absence in control experiments suggests that they are a direct result from PFA fixation, and not a sign of confocal/STED phototoxicity. Morphometric analysis of dendritic spines revealed that PFA application leads to wider spine necks, in line with previous findings, where spine necks were 30% thinner in cryogenically than chemically fixed samples (Tamada et al., 2020). While we did not see any changes in spine neck length, we observed slightly decreased spine head sizes. Overall, our “live-to-fixed-cell” experiments support the view that cryogenic protocols yield more trustworthy results than chemical fixation. Moreover, our study shows that fixation artifacts can occur in *ex vivo* preparations as a direct consequence of PFA immersion of the tissue, in addition to artifacts stemming from the *in vivo* response to transcardial perfusion.

PFA treatment (via incubation of slices or transcardial perfusion *in vivo*) is likely not the only culprit when it comes to fixation artifacts. Biological samples for microscopic analysis often undergo multiple preparatory steps, for instance dehydration, resin embedding or fixation with a combination of chemicals, such as glutaraldehyde or osmium tetroxide, to improve ultrastructural preservation and image contrast. These steps may also cause artifacts, such as electron dense granules (Hendriks and Eestermans, 1982), or organelle shrinkage (Mollenhauer, 1993). To avoid these artifacts, cryofixation involving high-pressure freezing was developed as an alternative approach, which preserves ECS shape and volume more faithfully (van Harreveld and Steiner, 1970). However, high-pressure freezing is technically more laborious, and has limited penetration depth, calling for ways to make chemical fixation less problematic, while retaining its accessibility and versatility.

To summarize, our time-lapse super-resolution approach enabled the direct comparison between live and fixed conditions within the same tissue sample at the nanoscale. Whereas short-lasting fixation (< 30 minutes) is largely innocuous to tissue nanostructure, longer PFA applications unmistakably lead to structural artifacts.

With the proliferation of super-resolution techniques relying on chemical fixation, it is crucial to reveal possible artifacts caused by chemical fixatives, ambient conditions (e.g. temperature) and other sample preparation steps, in order to optimize fixation protocols (Laporte et al., 2022; Pereira et al., 2019; Whelan and Bell, 2015).

Our new approach in combination with single-molecule based super-resolution techniques (Inavalli et al., 2019) to look at nanoscale morphology and protein arrangements may prove very useful for working out effective and practical solutions to increase the preservation of fixed cells and tissues and the fidelity of their microscopic analysis. Finally, our study also presents a case for the development and use of live-cell super-resolution microscopy, delivering data on the natural and dynamically evolving state of the biological system free of concerns of fixation artifacts of whatever provenance.

## Materials and Methods

### Mouse line

Animal experimental procedures were in accordance with the French National Code of Ethics on Animal Experimentation and approved by the Committee of Ethics of Bordeaux. All procedures were performed according to the guidelines of the European Directive 2010/63/UE.

Mice were housed under a 12 h light/12 h dark cycle at 20-22 °C with *ad libitum* access to food and water in the animal facility of the Interdisciplinary Institute for Neuroscience (University of Bordeaux/CNRS) and monitored daily by trained staff. All animals used were free of any disease or infection at the time of experiments. Pregnant females and females with litters were kept in cages with one male. We did not distinguish between males and females among the perinatal pups used for organotypic cultures, as potential anatomical and/or physiological differences between the two sexes were considered irrelevant in the context of this study.

C57Bl/6J wild-type mice were used for all experiments in this study.

### Organotypic brain slices

Organotypic hippocampal slices (Gähwiler, 1981) were dissected from 5 to 7 days old mice, and were cultured 2-5 weeks in a roller drum at 35 °C (for more details. See Tønnesen et al., 2018). Once a week, 500 µl of medium was exchanged in the tubes. For experiments, a given coverslip with a slice was mounted in an imaging chamber, and the slice was imaged from below through the glass coverslip, while it could be approached with PFA-containing solutions.

### Viral injections

In order to fluorescently label neurons or astrocytes, we have introduced either a Sindbis-Citrine or AAV2/1.gfaABC1D-Clover viruses to the brain slices via microinjections using a glass pipette connected to Picospritzer (Parker Hannifin). Briefly, the virus was injected via a pipette positioned into the CA1 area of the slice by brief pressure pulses (30 ms; 15 psi). For imaging of the neurons Sindbis-Citrine virus was injected into 2-weeks old wild-type slices 1 day prior to the experiments. To image astrocytes, 2-weeks old wild-type slices were injected with AAV2/1.gfaABC1D-Clover 2 weeks before the experiments.

### Extracellular labeling

Extracellular labeling of organotypic slices was performed as described before (Tønnesen et al., 2018). In brief, once the slice was transferred to the imaging chamber, it was immersed in 200 µM Calcein dye (Dojindo Laboratories) diluted in HEPES-based ACSF.

### Chemical fixation

After acquiring a live-image of either an extracellularly labeled or positively (neurons or astrocytes) labeled slice, the Calcein/ACSF or only ACSF solution was carefully extracted with a pipette to avoid any drift of the slice. Subsequently, a solution containing 4% PFA and 200 µM Calcein, both diluted in HEPES-based ACSF, or only 4% PFA diluted in HEPES-based ACSF, were pipetted on top of the slice. To minimize the evaporation of PFA, the imaging chamber was covered with a lid.

For overnight chemical fixation, the organotypic slices on a glass coverslip were transferred from the roller drum tube to a 6-well plate and instantly immersed in 4% PFA diluted in 1xPBS solution. The 6-well plate was placed at 4 °C overnight, followed by 3 washes in PBS. Finally, the fixed slices on the glass coverslip were mounted onto the imaging chamber containing Calcein/ ACSF solution.

### 3D-STED microscopy

We used a home-built 3D-STED setup (for details, see Inavalli et al., 2019) constructed around an inverted microscope body (DMI 6000 CS, Leica Microsystems), which was equipped with a TIRF oil objective (x100, 1.47 NA, HXC APO, Leica Microsystems) and a heating box (Cube and Box, Life Imaging Services) to maintain a stable temperature of 32°C. A pulsed-laser (PDL 800-D, PicoQuant) was used to deliver excitation pulses (90 ps at 80 MHz) at 485 nm and a synchronized de-excitation laser (Onefive Katana 06 HP, NKT Photonics) operating at 592 nm was used to generate the STED light pulses (500-700ps). The STED beam was reflected on a spatial light modulator (Easy3D Module, Abberior Instruments) to generate a mixture of doughnut- and bottle-shaped beams for 2D and 3D-STED respectively. Image acquisition was controlled by the Imspector software (Abberior Instruments). The performance and spatial resolution of the microscope was checked and optimized by visualizing and overlapping the PSFs of the laser beams using 150 nm gold nano-spheres and correcting the main optical aberrations. Usually, the spatial resolution was 175 nm (lateral) and 450 nm (axial) in confocal mode and 60 nm (lateral) and 160 nm (axial) in STED mode.

### Image acquisition

For imaging, slices were transferred on their glass coverslip to an imaging chamber and immersed in an imaging medium (artificial cerebrospinal fluid, ACSF) consisting of (in mM): 119 NaCl, 2.5 KCl, 1.3 MgSO_4_, 1 NaH_2_PO_4_ x 2H_2_O, 2.5 CaCl_2_ x 2H_2_O, 20 D-Glucose x H_2_O and 10 HEPES (all from Sigma Aldrich); 300 mOsm; pH 7.4. Confocal images were 100 × 100 × 4 µm^3^ z-stacks with a pixel size of 48.8 nm and Δz size of 1 µm. STED images were either 100 × 100 µm^2^ single plane acquisitions, 15 × 15 × 1 µm^3^ or 25 × 25 × 1 µm^3^ z-stacks with a pixel size of 19.53 nm, Δz size of 200 nm and a pixel dwell time of 30 µs. The excitation power was 0.5 µW and STED power was 30 mW at the entrance pupil of the objective.

### Image processing and statistical analysis

SUSHI images are single images taken from z-stacks or time-lapse series, as indicated. Images of astrocytes and dendrites are shown as maximum intensity z-projections. All morphometric measurements (widths or areas) of positively labeled structures were done on raw images in ImageJ (NIH), using the ‘Plot Line Profile’ function after drawing 3-pixel-wide straight lines across the structure of interest. Gaussian fits were applied directly in ImageJ and widths were calculated as FWHMs. Brightness and contrast were adjusted for each individual image and the look-up tables (LUT) were ‘grays’ for ECS and ‘orange hot’ for cellular structures. To calculate the volume fraction of the ECS, images were first binarized using a wavelet-based software, SpineJ (Levet et al., 2020) (**Fig. 1A**) and the fluorescence fraction was then calculated using ImageJ and expressed in percentage. Morphological parameters of dendritic spines were performed using the SpineJ software. The software identifies the neck region and places lines that are orthogonal to the neck axis at regular of 75 nm. It then calculates the FWHM of the neck diameter, returning the minimum, maximum and median values for each analyzed spine. We limited the morphology analysis to dendritic spines with clear neck and head compartments, commonly referred to as mushroom spines.

Statistical tests were performed using Graphpad Prism software. Normally distributed data are presented as mean with standard deviation, while non-normal data are presented as median with interquartile range. The size and type of individual samples, n, for given experiments is indicated and specified in the results section and in figure legends. Asterisks in figures indicate p values as follows: * p < 0.05, ** p < 0.01, *** p < 0.001, **** p< 0.0001.

## Acknowledgements

The authors thank R. Sterling for preparing PFA solutions, constructing sample holder and ensuring safety measurements for handling PFA solutions. We thank F. Quici, J. Angibaud and IINS Cell Culture Facility for support with organotypic slice cultures. We thank A. Boyce and Y. Dembitskaya for comments on the manuscript.

## Author contributions

AI carried out all experiments and analysis. MA provided student supervision, technical and intellectual input. VVGKI and SB provided technical support for the STED microscopy. UVN conceived the study and provided supervision. The paper was written by AI, MA and UVN with input from all authors.

